# Individuals with 22q11.2 deletion syndrome show intact prediction but reduced adaptation in responses to repeated sounds: evidence from Bayesian mapping

**DOI:** 10.1101/441634

**Authors:** Kit Melissa Larsen, Morten Mørup, Michelle Rosgaard Birknow, Elvira Fischer, Line Olsen, Michael Didriksen, William Frans Christiaan Baaré, Thomas Mears Werge, Marta Isabel Garrido, Hartwig Roman Siebner

**Affiliations:** Danish Research Centre for Magnetic Resonance, Centre for Functional and Diagnostic Imaging and Research, Copenhagen University Hospital Hvidovre, Hvidovre, Denmark; DTU Compute, Cognitive Systems, Technical University of Denmark, Lyngby, Denmark; Institute of Biological Psychiatry, Mental Health Centre Sct. Hans, Copenhagen University Hospital, Boserupvej 2, DK-4000 Roskilde, Denmark; iPSYCH, The Lundbeck Foundation Initiative for Integrative Psychiatric Research, Aarhus and Copenhagen, Denmark; Queensland Brain Institute, The University of Queensland, St Lucia 4072, Brisbane, Australia; Synaptic Transmission, H. Lundbeck A/S, Ottiliavej 9, DK-2500 Valby, Denmark; Centre for Advanced Imaging, The University of Queensland, St Lucia 4072, Brisbane, Australia; Australian Research Council Centre of Excellence for Integrative Brain Function Centre of Excellence for Integrative Brain Function, The University of Queensland, St Lucia 4072, Brisbane, Australia; School of Mathematics and Physics, The University of Queensland, St Lucia 4072, Brisbane, Australia; Department of Clinical Medicine, Faculty of Health and Medical Sciences, University of Copenhagen, Copenhagen, Denmark; Department of Neurology, Copenhagen University Hospital Bispebjerg, Copenhagen, Denmark

**Keywords:** 22q11 Deletion Syndrome, Dynamic causal modelling, Posterior probability maps, EEG, Mismatch negativity, repetition suppression

## Abstract

**Background:** One of the most common copy number variants, the 22q11.2 microdeletion, confers an increased risk for schizophrenia. Since schizophrenia has been associated with an aberrant neural response to repeated stimuli through both reduced adaptation and prediction, we here hypothesized that this may also be the case in nonpsychotic individuals with a 22q11.2 deletion.

**Methods:** We recorded high-density EEG from 19 individuals with 22q11.2 deletion syndrome (12-25 years), as well as 27 healthy volunteers with comparable age and sex distribution, while they listened to a sequence of sounds arranged in a roving oddball paradigm. Using posterior probability maps and dynamic causal modelling we tested three different models accounting for repetition dependent changes in cortical responses as well as in effective connectivity; namely an adaptation model, a prediction model, and a model including both adaptation and prediction.

**Results:** Repetition-dependent changes were parametrically modulated by a combination of adaptation and prediction and were apparent in both cortical responses and in the underlying effective connectivity. This effect was reduced in individuals with a 22q11.2 deletion and was negatively correlated with negative symptom severity. Follow-up analysis showed that the reduced effect of the combined adaptation and prediction model seen in individuals with 22q11.2 deletion was driven by reduced adaptation rather than prediction failure.

**Conclusions:** Our findings suggest that adaptation is reduced in individuals with a 22q11.2 deletion, which can be interpreted in light of the framework of predictive coding as a failure to suppress prediction errors.

## Introduction

22q11.2 deletion syndrome (22q11.2DS) is caused by one the most common copy number variants in humans with a prevalence of 1:2000 to 1:4000^1–4^. The 22q11.2DS is clinically presented with a highly variable phenotype, including a range of somatic disorders, learning problems, cognitive deficits^5,6^. 22q11.2DS is associated with a high frequency of several neurodevelopmental disorders, including autism spectrum disorder, attention deficit hyperactivity disorder and schizophrenia^6–11^, with the prevalence of schizophrenia-spectrum disorder being estimated at 41% in adult individuals with 22q11.2DS^9^.

The ability to adapt to the ever changing environment and react to deviations within it, is something the healthy brain masters on a daily basis. However, people with schizophrenia show reduced ability to adapt to the environment expressed as a state of aberrant salience^12^. A neurophysiological example of this phenomenon is the typically reduced neural response to repeated stimuli, a process called repetition suppression (RS), often depicted as a consequence of neural fatigue^13^. Recent theoretical formulations inspired on predictive coding^14–16^ propose that altered RS in schizophrenia may be caused by inaccurate sensory predictions^17–20^. According to this perspective, RS is a consequence of prediction error minimization afforded by adaptation to the environment through learning about incoming sensory input. Repetition-dependent changes in responses to repeated stimuli have previously been explained by experience-dependent changes in effective connectivity both in extrinsic connections, between brain areas, encoding predictions as well as intrinsic, within brain area, connections believed to encode prediction precision^21^. Most event-related potential (ERP) studies have focused on the first repetition relative to the initial presentation^22–25^. Very recent, Stefanics and colleagues^23^ showed that RS was best explained by an exponential model indicating that repetition effects are observable for trials beyond the first repetition, highlighting the necessity to investigate brain responses beyond the first repetition in order to understand the underlying processes of RS.

RS in schizophrenia has mostly been studied through sensory gating where suppression of P50 is seen to be reduced^26,27^. However, RS has also been shown to be intact in schizophrenia^28^, manifest in comparable ERPs to repeated auditory tones^29^. RS is sparsely studied in 22q11.2DS with again opposing results where sensory gating as indexed by P50 has been shown to be sometimes intact^30,31^ and other times impaired^32,33^. Given that the underlying mechanism of RS in 22q11.2DS is still poorly understood, we investigated the brain mechanism underpinning RS in 22q11.2DS and how these mechanisms might potentially deviate from what is seen in healthy controls. We formalized three theoretical models to explain RS: the adaptation model, the prediction model and finally the combined model, in the following referred to adaptation&prediction. These three models were tested both at the scalp level using Bayesian mapping for M/EEG^34–36^ and at the connectivity level using dynamical causal modelling (DCM)^37^. Firstly, we hypothesized that responses to repeated stimuli would show a parametric modulation with an overall decrease in connectivity within the tested network for the first repetitions, followed by an increase reflecting the prediction of new stimuli, in agreement with the combined adaptation&prediction model. Next, we investigated group differences in 22q11.2DS and healthy controls at the scalp level and connectivity level within the model that best described RS. This unique way of modelling RS allowed us to pinpoint the origin of potential deficits in 22q11.2DS, namely the adaptive and predictive processes underpinning RS.

## Materials and Methods

Participants and stimuli administered here have been previously described^38^, where we report on mismatch negativity (MMN) responses in 22q11.2DS. For the purpose of clarity and completion we provide a summary below.

### Participants

This study is part of a larger Danish nationwide initiative^39^. A group of 19 non-psychotic individuals with a verified deletion of 3Mbs at chromosome 22q11.2 with no current or previous diagnosis were included. 27 healthy individuals without 22q11.2DS was included as a control group with comparable age distribution (controls age range: 12-25 years; mean age: 15.96, standard deviation (SD) = 2.71 years; 22q11.2DS mean age: 15.47, SD 2.41 years, t_44_=-0.63 p = 0.53) and sex ratio (male/female controls: 18/9, cases: 13/6, χ2 = 0.02, p = 0.90). The following exclusion criteria were applied to controls: presence of a) schizophrenia, schizotypal and delusional disorders (ICD10 DF20- 29); b) bipolar disorder (ICD10 DF30-31); c) depression (ICD DF32-33) except for a past episode of mild or moderate depression (ICD10 DF 32.0 or 32.1); d) substance abuse; or e) a first degree relative with a psychotic illness. Screening for current psychosis and rating the severity of schizophrenia-related symptoms was done using the Structured Interview for Prodromal Syndromes ^40,41^. Schizophrenia-related symptoms were assessed within the four domains: positive, negative, disorganized and general symptoms. The regional Ethical Committee of Copenhagen (project id: H-3-2012-136) and the Danish Data protection Agency (project id: 2007-58-0015) approved the study. All participants underwent a verbal and written informed consent process. Participants under the age of 18 provided a verbal assent while their parent’s completed written consent. For a thorough description of clinical tests and demographics for the included participants, please see supplementary material.

### Stimuli

The roving paradigm was adapted from^42^ and comprised of roving sequences of sounds ranging from 1 to 8 drawn from a discrete uniform distribution, **Error! Reference source not found.**A. With this paradigm it is possible to study the responses to repeated stimulation and thereby the parametric effect of repetition. Participants sat in a comfortable chair and watched a silent movie displaying underwater scenery free of any sudden or salient visual events during the 15 minutes of recording. Participants were instructed to ignore the sounds. Audiometric testing was performed prior to the experiment, see supplementary material.

### Data acquisition and pre-processing

EEG data were recorded using a 128 channel ActiveTwo Biosemi System (BioSemi, Amsterdam, Netherlands), with a sampling frequency of 4096 Hz. Pre-processing included; band pass filtering between 0.5Hz - 40Hz using a second order Butterworth filter, downsampling to 500Hz and finally epoching with a peristimulus window of −100ms to 400ms. The preprocessing was carried out using EEGLAB (Delorme and Makeig, 2004). Baseline correction was applied using the average over the time window −100ms to −10ms. Re-referencing to the average reference, artefact removal, scalp analysis, and the DCM analysis were performed using SPM12 (http://www.fil.ion.ucl.ac.uk/spm/). Epochs were rejected if their values exceeded ±100 μV. One of the participants (belonging to the 22q11.2DS group) was discarded because the majority of epochs were rejected with this approach (above 80%).

### The three models accounting for repetition dependent effects

Since we were interested in the repetition dependent changes in ERPs and effective connectivity, we explored three different models, given below for tone *r* = 1,…,9.

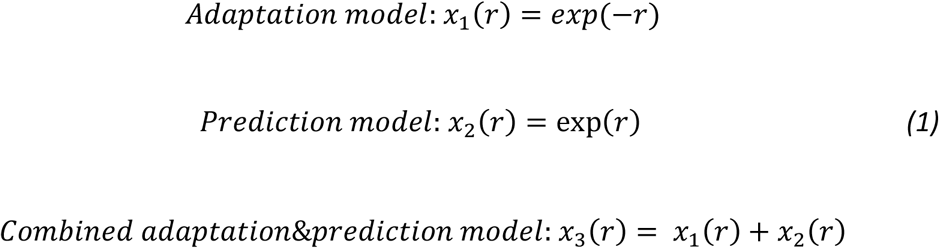

We chose the exponential function, given that responses are typically seen heavily reduced in the first repetition, whereas responses seem to become similar thereafter, which is in line with recent findings^23^, see Figure 1B. The adaptation model postulates that responses decrease with the number of repetitions. Conversely, the prediction model postulates that responses will increase with repetitions, reflecting formation of an expectation that a new event will occur. Finally the adaptation&prediction model is a combination of the adaptation and prediction model in that the initial exponential decay will capture changes due to habituation or adaptation and the growing exponential towards the end will capture formation of an expectation, or prediction.

**Figure 1:**
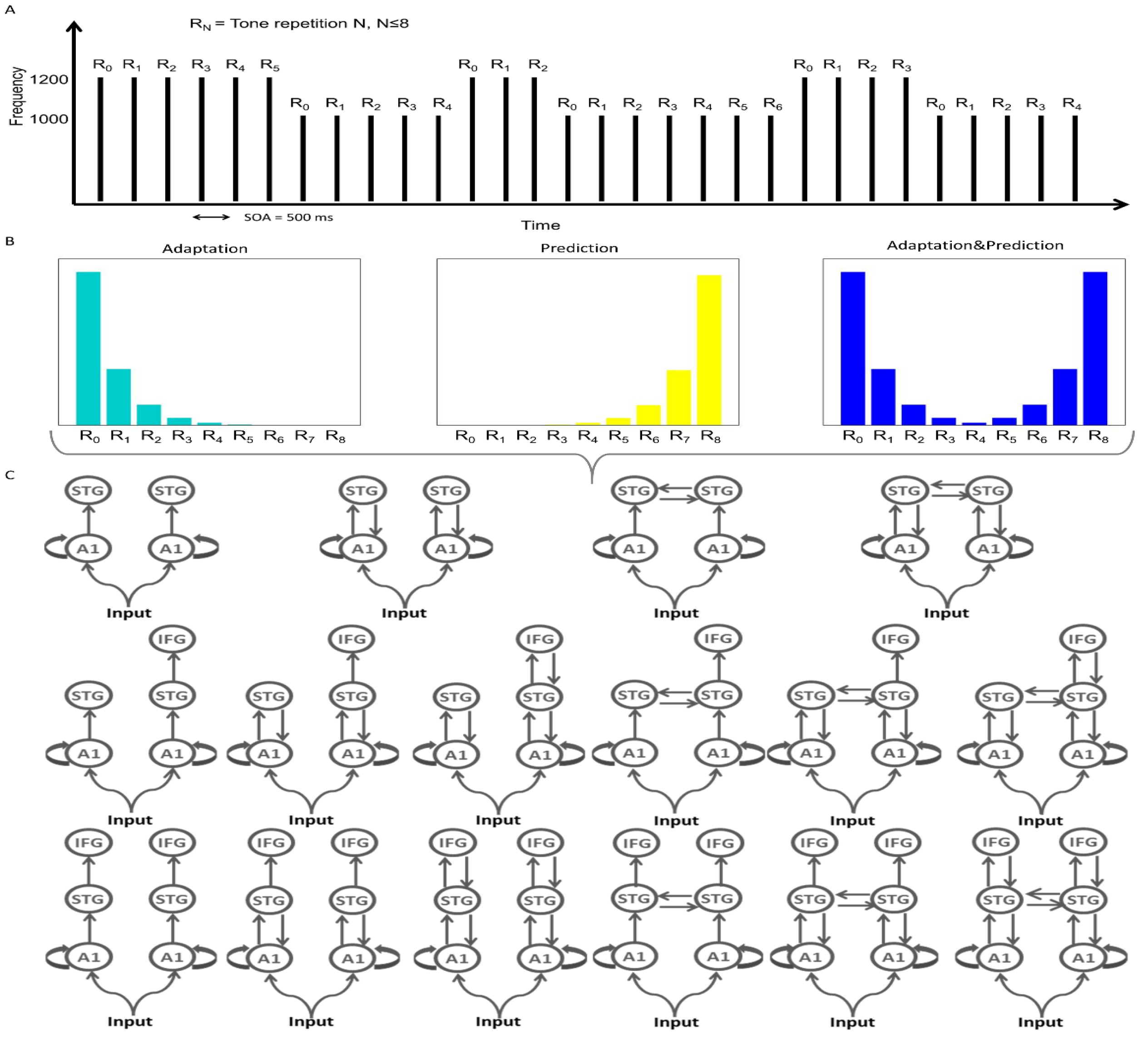
Experimental design of the roving paradigm and the three different repetition effects models. A: The tone repetition, R_N_, varies randomly between 0 and 8 (maximum of 9 tones). The sequences of tones vary by having a frequency of either 1000 Hz or 1200 Hz. Stimulus onset asynchrony is fixed at 500ms. B: The three parametric models for repetition-specific effects: the adaptation model, the prediction model and the adaptation&prediction model. C: Model space for DCM models. Each family consisted of the same DCMs, but deviates in the parametric modulation between conditions, that is, the effect that repetitions of tones has on ERPs.

**Figure 2:**
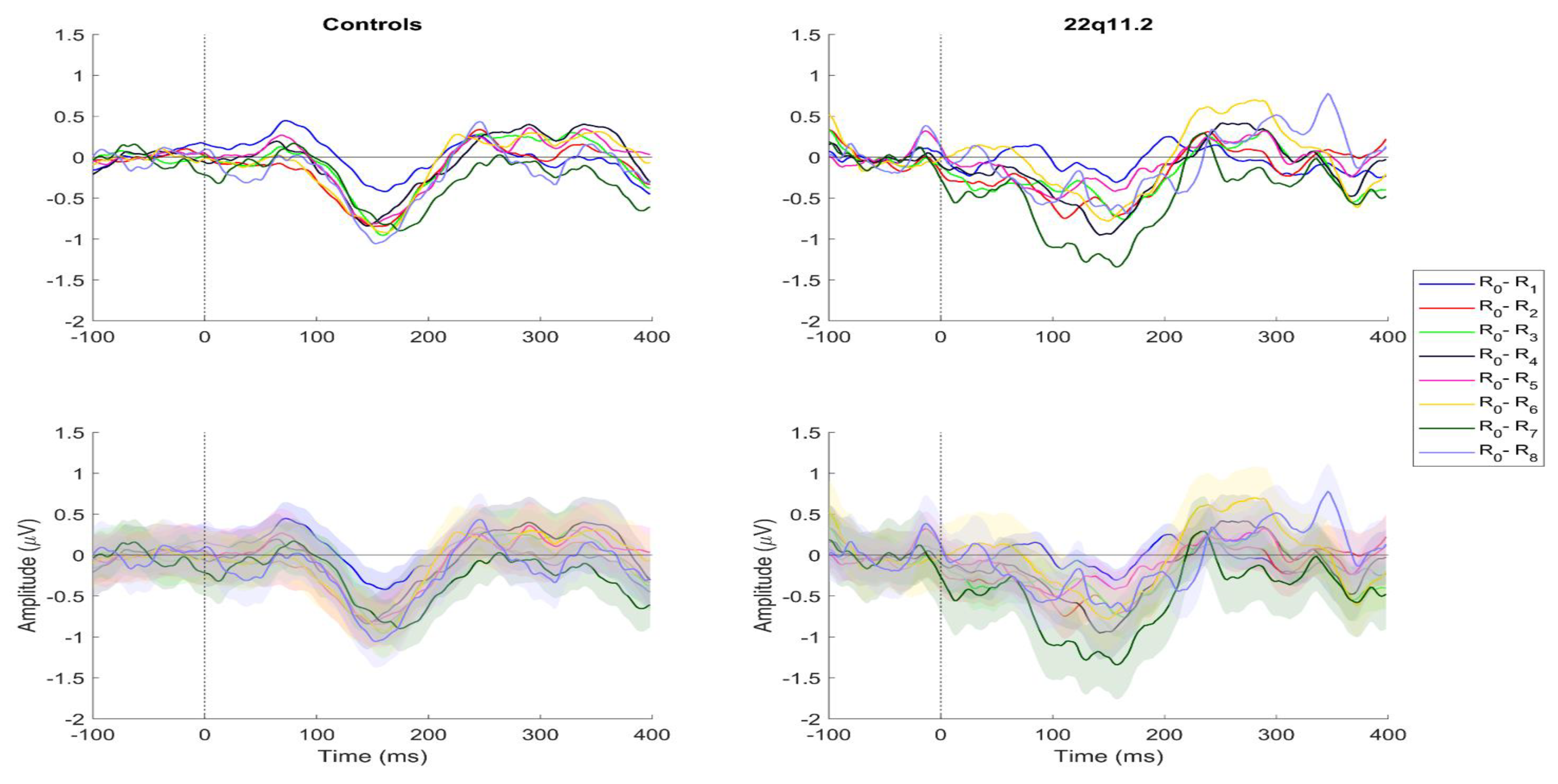
Grand average difference responses for controls and 22q11.2DS from channel Fz. Left: responses to each tone repetition for controls. Right: Corresponding responses for individuals with 22q11.2DS. First row represent the mean of the responses whereas the second row represents the mean with the shaded area representing one standard deviation from the mean.

### Posterior Probability Maps

To compare the adaptation, prediction, and adaptation&prediction models for ERPs, we used posterior probability maps^34–36^. Epoched data were converted into scalp-map images of dimension 64×64 obtained using interpolation. After the conversion to scalp-map images smoothing using a Gaussian kernel specified by a FWHM of 8 mm^2^ in the spatial dimension and 10ms in the temporal dimension was performed.

Individual participant voxel-wise whole brain log-evidences were calculated using regressors describing the hypothesized relationship amongst the tone repetitions i.e. the equations in (1). The log-evidence for each model were estimated using the variational Bayes first-level model specification^43^. Group level (pooled across controls and 22q11.2DS) posterior probability maps were calculated using the random effects approach (RFX) for each model. These probability maps can then be used to compare between the three different models for each voxel and time point.

### Dynamic Causal Model specification

To investigate the underlying connectivity network of RS, we used DCM which is a hypothesis driven method for estimating effective connectivity between brain areas^37,44^. We have previously used the same paradigm to study MMN responses in 22q11.2DS^38^ where we formulated a set of models motivated by previous studies on MMN generators^45–48^ as well as previous model comparisons of MMN generation^42,49^. This network includes bilateral sources in the primary auditory cortex (A1), superior temporal gyrus (STG) and inferior frontal gyrus (IFG), with the IFG usually being most consistent in the right side. The bilateral sources in A1 receive the input. RS in a roving MMN paradigm has been previously studied using DCM^21^ with bilateral A1 and STG sources being included. We defined 16 models starting with the right and left A1 and building up the remaining models by adding hierarchical levels until we had a full network comprising the six sources: bilateral A1, STG and IFG, **Error! Reference source not found.**B. Each of the parametric DCM models were estimated for each participant individually with all nine tones in the same model. We defined each of these parametric forms as families that only deviate in the specific condition-specific parametric effect.

### Bayesian Model Selection

RFX Bayesian model selection was used for the pooled group to test which of the repetition dependent models best described the data overall both at the scalp level from the posterior probability maps as well as from the DCMs. Posterior and exceedance probabilities were used to compare the models.

### Assessing group differences at the scalp level and connectivity level

#### Spatiotemporal analysis

In order to test for group differences within the winning model at the scalp level, spatio-temporal analysis was performed over the whole sensor-space (i.e. all electrodes) and time (0 ms to 400 ms) using a full factorial 2×9 design with factors group (controls and 22q11.2DS) and condition (repetitions) and age and sex included as covariates. Weights under the winning model, given by equation (1) were entered as contrast allowing to asses group differences in the parametric effect present i.e., how much of the winning model is present in controls compared to 22q11.2DS. All p values reported are thresholded using p<0.05 FWE corrected at cluster level. To enable the investigation as to whether scalp data activity was correlated with the clinical symptoms in the 22q11.2DS group, we extracted activity from single-participant contrast images associated with the parametric effect. The activity was extracted from a square region with a size of 10mm x 10 mm around the peak difference between controls and patients with respect to the winning model at the scalp level. Correlation with negative symptoms were performed using Pearson correlation. Only correlations with negative symptoms were done since the occurrence of positive, disorganized and generalized symptoms was very low and skewed in the 22q11.2DS group, see demographics in the supplementary material.

#### Connectivity parameters

Bayesian model averaging (BMA) was carried out within the family with highest exceedance probability, to allow for group comparison of the connectivity parameters (B-parameters)^50^. A one-way ANCOVA with group as a factor (controls and 22q11.2DS) and age and sex as covariates was performed for each of the parameters. Results are reported both uncorrected as well as corrected for multiple comparison using Bonferroni.

## Results

### Repetition dependent changes can be explained by a combination of adaptation and prediction both in cortical responses and in effective connectivity

**Error! Reference source not found.** shows difference waves for each repetition of the tones for controls and individuals with 22q11.2DS. Visually it can be seen that controls (left column) show a clear jump from first repetition (in blue) compared to the rest of the repetitions whereas there seems to be no clear pattern between responses to the tone repetitions in the 22q11.2DS group (right column).

At the scalp level the posterior probability maps showed that the combined adaptation&prediction model outperformed the adaptation and prediction models throughout all time points, see **Error! Reference source not found.**A and B, where probabilities are shown summed across space and pooled across the two groups (controls and 22q11.2DS). However, the model’s outperformance was most pronounced in the middle of the epoch, from 50ms to 350ms. Summing across both time and space (**Error! Reference source not found.**C), the model probability for the combined model outperforms the two other models. It is observed that the model probabilities in the baseline period (Figure 3E) is close to equal meaning that the bias towards a winning model is very small. The exceedance probability of the DCMs for the combined model is 1 (Figure 3D), meaning that at the connectivity level, this model is a clear winner as well. Together, these results show that of the considered hypotheses repetition-dependent changes in ERPs and in effective connectivity are best explained by adaptation and prediction formation. The spatial distribution of the probabilities in Figure 3F, thresholded at posterior probability p = 0.83, shows that the spatial distribution of the combined adaptation & prediction model involves electrodes throughout the fronto-central area.

**Figure 3:**
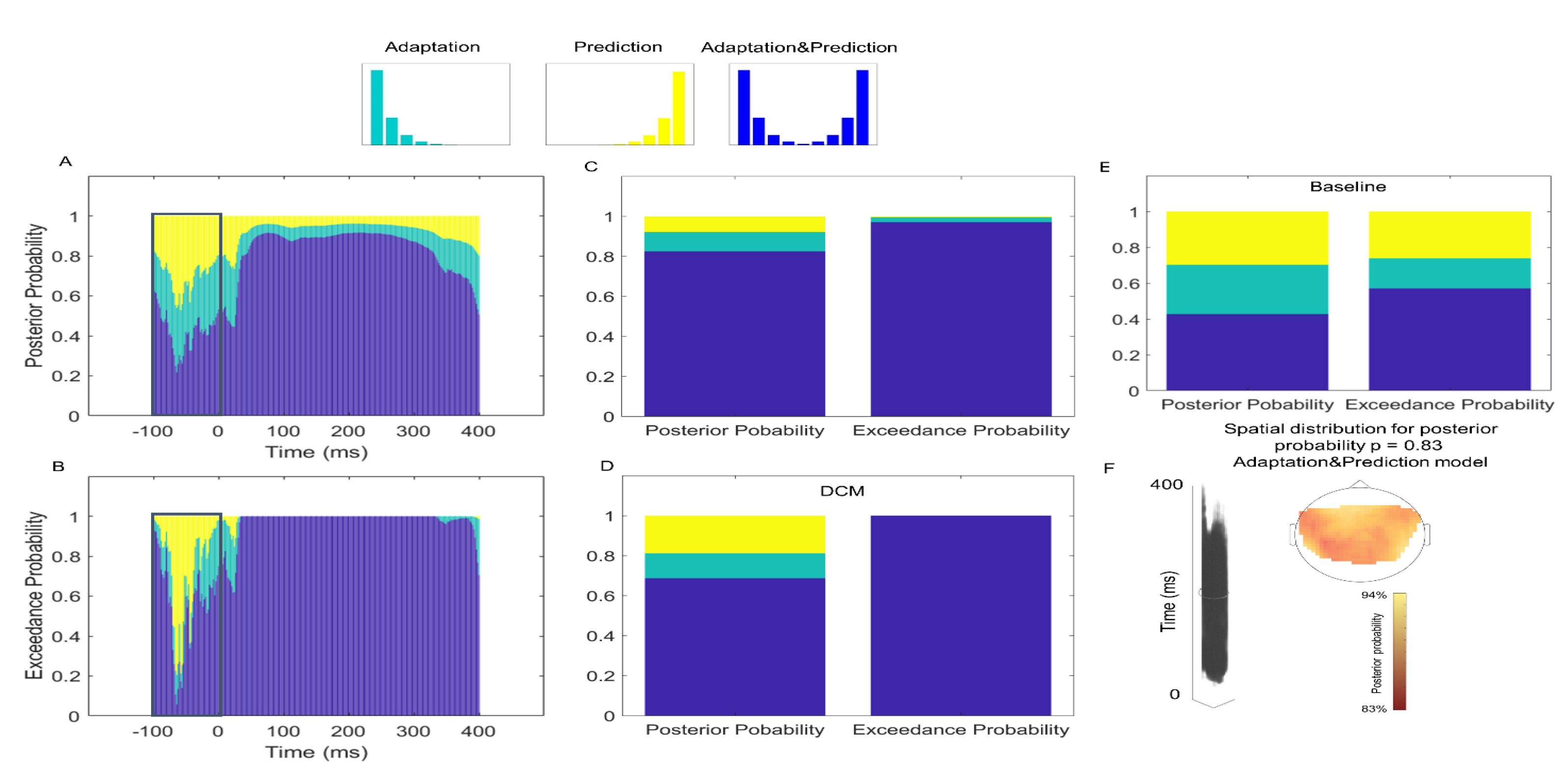
Exceedance and posterior probabilities of the three models. A: posterior probability as a function of time, summed across space for the adaptation model (turquoise), the prediction model (yellow) and the combined adaptation&prediction model (blue). B: Same as A, for exceedance probabilities. C: Posterior and exceedance probability for the model comparison at the scalp level, summed across space and time. D: Posterior and exceedance probability for DCMs with connectivity modulations according to the adaptation, prediction, and the adaptation&prediction family. E: Same as C, for the baseline period only. F: Spatial distribution of the combined adaptation&prediction model thresholded at posterior probability p = 0.83.

**Figure 4:**
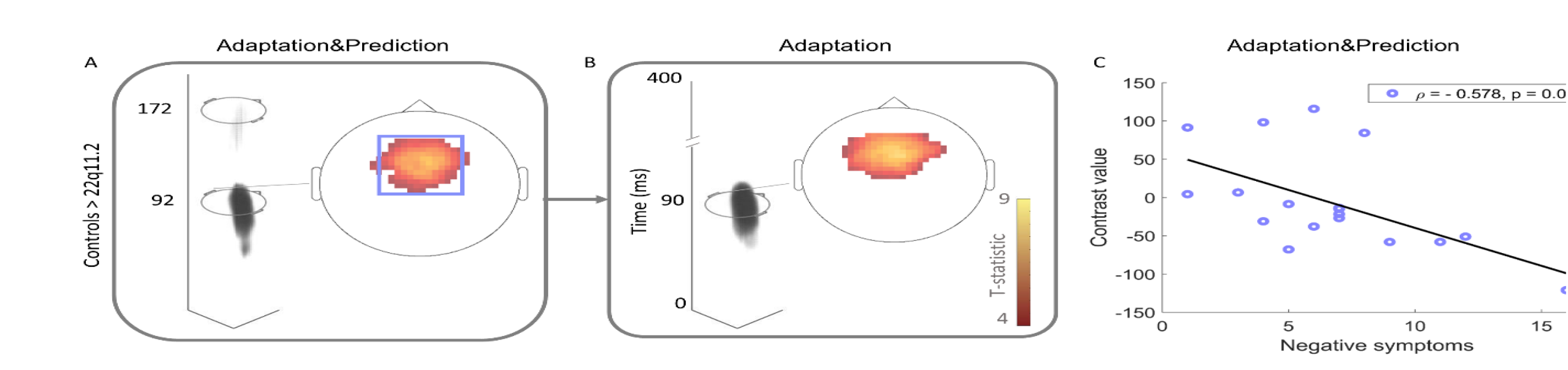
T-maps show that reduced repetition suppression in individuals with 22q11.2DS is driven by attenuated adaptation processes. A: Group effect for the winning adaptation&prediction model. There is a significant cluster in the fronto-central area peaking at 92 ms. B: To determine the driver of this effect, we also show the map of the main effect of group for the adaptation model (no effect observed for the prediction model). All results are shown at p < 0.05 FWE corrected at cluster level. C: Correlation with negative symptoms. The activity extracted from the peak of the controls > 22q11.2 contrast for the adaptation&prediction model. The higher degree of negative symptoms in 22q11.2DS were associated with less amount of activation for the controls greater than 22q11.2 adaptation&prediction contrast.

Within the combined adaptation&prediction model the DCM model selection of the models shown in Figure 1C, did not reveal a clear winning model, which is why we kept the model comparison at the family level.

### 22q11.2DS show reduced adaptation but not prediction

Knowing that repetition dependent changes are explained by a combination of adaptation and prediction processes manifested both in the ERPs and in condition-dependent connectivity changes, we next tested for group differences within this model. Analysis of the scalp maps of the repeated stimuli for the contrast controls greater than 22q11.2DS for the combined adaptation&prediction model revealed an effect peaking at 92 ms (see **Error! Reference source not found.**). Further, we observed an effect around 172 ms whereas the reversed contrast, 22q11.2DS greater than controls revealed an effect at 74 ms. However, the effect at 74ms was a small set of points that fell in the interpolated area where no electrodes were positioned. We therefore see this effect as spurious and have not depicted it in **Error! Reference source not found.**. To delineate if the group difference at 92 ms was driven by the adaptation or the prediction processes, we looked at the group effects for the adaptation model and prediction model (**Error! Reference source not found.**B). Our results indicate that the group difference was driven by the adaptation process, suggesting that the ability to adapt to the tones is impaired in individuals with 22q11.2DS.

We then investigated whether the degree to which repetition dependent changes in cortical responses in 22q11.2DS were associated with symptomatology. To do so, we restricted our search to a square region round the peak difference in the controls greater than 22q11.2 adaptation&prediction contrast. The strength of the parametric modulator extracted from this region for each individual with 22q11.2DS was negatively correlated with the severity of negative symptoms (**Error! Reference source not found.**C)(ρ = −0.578, p = 0.012, unc., p = 0.024 corrected). Hence, the more severe the negative symptoms, the less activity in the areas associated with the adaptation&prediction model.

Since 22q11.2DS is associated with hearing loss^51^ and lower IQ levels^52,53^ we did a post-hoc analysis to test whether these variables could explain the observed effects in the spatio-temporal analysis. An effect of IQ was found in fronto-temporal channels at around 150 ms whereas an effect of hearing levels were found at central electrodes around 350 ms. However, the group effects persisted even after adding hearing levels and IQ as a covariate in the analysis, suggesting that even though these showed effects on the EEG, they do not fully account for the observed group effects.

### Group differences in connectivity strength

Individuals with a 22q11.2 deletion showed a stronger modulation than controls in the extrinsic connection from right IFG to right STG (F_1,41_ = 6.147, p = 0.017) in the B parameter associated with an adaptation effect. However, it did not survive correction for multiple comparisons using a conservative Bonferroni correction for 12 test, i.e. connections (α = 0.05/12 = 0.004). There was no group difference observed in the B parameters associated with the prediction effect. There was no effect of the covariates sex and age. The group effect persisted when adding hearing levels as covariate, but disappeared when adding IQ as a covariate, suggesting that IQ was driving this effect.

## Discussion

This study provides evidence that adaptation to repeated sounds is diminished in a group of young non psychotic individuals with 22q11.2DS. Our results suggest that repetition-dependent changes both in ERPs and in effective connectivity are modulated by a combination of adaptation and prediction processes. Furthermore, we found that group differences in the relationship between ERP activity and stimuli repetition was driven by reduced adaptation in individuals with 22q11.2DS in the early ERP component N1, at fronto-central electrodes. Critically, the degree to which this relationship between ERP activity over repetitions was present, correlated negatively with the degree of negative symptoms in 22q11.2DS. Results therefore suggest that repetition dependent changes in cortical responses in 22q11.2DS are associated with negative symptoms.

RS is characterized by a reduction in neural activity, or adaptation, caused by repeated stimuli^54–57^, a phenomenon thought to be mediated by synaptic communication. Predictive coding theories have re-interpreted RS as the neural mechanism underpinning perceptual learning and inference^14,15^. Our findings suggest that repetition-dependent changes both in ERPs and in brain connectivity can be explained by a model combining both adaptation and prediction processes. The adaptation component of our combined model predicts an initial exuberant prediction error occurring immediately after a change in sound statistics, which is then followed by decreases in neuronal responses caused by sound repetition. While this accounts for the neurophysiological data evoked by the first half of the sound trains, the prediction component resembles an expectation build-up, as if the participant began to expect an eventual change sometime during the second half of the sound trains. This is in line with the notion that responses to repeated sounds are not only caused by simple mechanisms as neural fatigue, but are likely to be caused by fulfilled expectations^15,17,18^ including forward message passing of prediction error and backward message passing of predictions or expectations. The spatio-temporal analysis within the combined adaptation&prediction model revealed a group difference driven by the adaptation component, whereas no difference was seen in the prediction component. This is in line with our previous work, where no difference was seen in MMN responses between the two groups (indicating that prediction is preserved)^38^. There is however opposing results in the literature on the change detection mechanism in 22q11.2DS. While MMN evoked by a duration deviant has been shown to be reduced in 22q11.2DS^58^, no difference across five deviants types; duration, frequency, gap, intensity and location was observed in^32^. While MMN was found to be preserved in^38^, we found a general increased response to tones, evidenced by increased N1 responses in 22q11.2, suggesting either increased sensitivity to tones, or reduced adaptation. Here, we show that the adaptation component is reduced in 22q11.2DS.

RS is very sparsely studied in 22q11.2DS with opposite results on sensory gating as well, with P50 shown to be sometimes intact^30,31^ and other times impaired^32^. However, sensory gating usually entails a paired-click paradigm, excluding the possibility of studying effects beyond the first repetition which have been shown to occur^23^. It is therefore hard to compare results from the present study to previous findings on P50 sensory gating.

In conclusion, we show that young non-psychotic individuals with 22q11.2DS are impaired at modulating neural activity to the environmental statistics associated with repeated stimuli.

## Acknowledgements

We thank all participants and families for taking their time to participate in our study and the Danish National 22q11DS Association for their strong support of our work. We highly appreciate the efforts of Anders Vangkilde and Henriette Schmock for their dedicated assistance in the recruitment and clinical assesments. We thank staff involved in the Danish Blood Donor Study, Capital Region Blood bank, Glostrup and http://www.forsogsperson.dk/ from which our control participants were recruited.

This study was funded by the Lundbeck Foundation, Denmark (R155-2014-1724); Lundbeck Foundation [Grant of Excellence “ContAct” R59 A5399]; Lundbeck Foundation fellowship (R105-9813); The Capital Region’s Research Foundation for Mental Health Research; the Australian Research Council Centre of Excellence for Integrative Brain Function (ARC Centre Grant CE140100007); University of Queensland Fellowship (2016000071) to MIG. H.R.S. holds a professorship in precision medicine at the Faculty of Health Sciences and Medicine, University of Copenhagen, sponsored by the Lundbeckfonden.

## Financial disclosures

H.R.S. received honoraria as speaker from Lundbeck A/S, Valby, Denmark, Biogen Idec, Denmark A/S, Genzyme, Denmark and MerckSerono, Denmark, honoraria as editor from Elsevier Publishers, Amsterdam, The Netherlands and Springer Publishing, Stuttgart, Germany, travel support from MagVenture, Denmark, and grant support from Biogen Idec, Denmark A/S. M.R.B is a prior employee at H. Lundbeck A/S, Denmark and received financial support for her PhD from the Innovation Fund Denmark. M.D is employed with Lundbeck A/S. The authors declare no further biomedical financial interests or potential conflicts of interest.

This article is distributed under the Danish legislation governed by the Privacy Act (act# 429, 31/05/2000), which does not permit data sharing at publicly available repositories or in raw formats. Summary statistics can be obtained through contact with the corresponding author.

## Abbreviations

DCM: Dynamic Causal Modelling
MMN: Mismatch Negativity
D;22q11.2DS: 22q11.2 Deletion Syndrome

